# Potential mechanisms of action of celastrol against rheumatoid arthritis: transcriptomic and proteomic analysis

**DOI:** 10.1101/2020.05.14.095737

**Authors:** Xinqiang Song, Erqin Dai, Yu Zhang, Hongtao Du, Lei Wang, Ningning Yang

## Abstract

**Background:** The natural triterpene celastrol exhibits potential anti-inflammatory activity in inflammatory diseases such as rheumatoid arthritis (RA).

**Methods:** Here we explored through what proteins and processes celastrol may act in activated fibroblast-like synoviocytes (FLS) from RA patients. Differential expression of genes and proteins after celastrol treatment of FLS was examined using RNA sequencing, label-free relatively quantitative proteomics and molecular docking.

**Results:** Expression of 26,565 genes and 3,372 proteins was analyzed. Celastrol was associated with significant changes in genes that respond to oxidative stress and oxygen levels, as well as genes that stabilize or synthesize components of the extracellular matrix.

**Conclusions:** These results identify several potential mechanisms through which celastrol may inhibit inflammation in RA.

## 1. Introduction

Rheumatoid arthritis (RA) is a chronic inflammatory disorder that affects multiple peripheral joints [1-3]. It is characterized by synovial hyperplasia, which results in joint destruction. One of the main cell types involved in RA pathology are fibroblast-like synoviocytes (FLS) [4, 5]. During RA progression, FLS grow in an anchorage-independent fashion, their morphology changes, and they express high levels of biomarkers such as c-fos, Jun-B, egr-1 and matrix metalloprotease (MMP)-3 [6]. FLS also secretes several inflammatory cytokines, such as IL-6, IL-8, IL-1, TNF-α and MCP-1 [7, 8]. How FLS are activated to drive RA is unclear, and epigenetics may play a role[9, 10].

The natural triterpene celastrol from *Tripterygium wilfordii* Hook f [11-14], is widely used in China to treat RA as well as autoimmune and inflammatory diseases [15-17]. Although the clinical efficacy of celastrol has been well-documented, its mechanism of action remains unclear. Studies *in vitro* have shown that high doses of celastrol can induce apoptosis and inhibit growth and invasion of FLS in RA [4, 7, 18]. However, we are unaware of comprehensive screens to identify FLS proteins and processes affected by celastrol.

High-throughput profiling of transcripts and proteins can identify genes whose expression is altered by a drug of interest [19-21]. Transcriptomic profiling on its own does not take into account post-transcriptional modifications, efficiency of translation or transcript turnover, all of which can affect activity of the encoded proteins[22, 23]. Therefore combining transcriptomic and proteomic profiling may be more effective for identifying drug-induced changes [24-26].

In the present study, we took advantage of technological advances in measuring mRNA levels by RNA sequencing (RNA-Seq) and protein levels by label-free relatively quantitative proteomics [27-30] in order to explore the effects of celastrol on FLS from RA patients.

## 2. Materials and Methods

### 2.1 Cell culture

Human FLS (as control), RA patient FLS and synoviocyte growth medium were purchased from Cell Applications (Santiago, California, USA). FLS were cultured at 37 °C in 5% CO_2_ in synoviocyte growth medium containing 100 μg/mL streptomycin and 100 U/mL penicillin. FLS in the exponential phase of growth were seeded into 10-cm dishes (1 × 10^6^ cells/dish).

After FLS had spread across the dishes, they were fixed for 15 min in 2% paraformaldehyde, blocked for 1 h with rabbit serum (Sigma), then incubated for 1 h with antibody against the FLS marker vimentin (1:50; Abcam, Cambridge, MA, USA). The dishes were washed three times with 0.01% saponin in phosphate-buffered saline (PBS) for 15 min each, then incubated for 1 h with secondary antibody conjugated with fluorescein (Jackson Immuno Research, West Grove, PA, USA). Dishes were washed in PBS, the nuclear stain DAPI was added, the coverslips were washed three times with 0.01% saponin in PBS for 15 min each, and then they were washed twice in PBS for 10 min each. The dishes were mounted on slides using anti-fade mounting medium and analyzed under an IX2-ILL100 fluorescence microscope (Olympus, Tokyo, Japan).

### 2.2 RNA extraction

Celastrol was dissolved in DMSO to make a 100 mM stock solution, which was diluted to the desired concentrations, such that the final proportion of DMSO in culture wells was 1%. As controls, Human FLS were treated with DMSO such that the final proportion in wells was 1%. FLS were seeded in medium, then incubated for 24 h with 5 μM celastrol (Shunbo Biotech, Shanghai, China) [4]. Total RNA was isolated from FLS using TRIzol® (Invitrogen) according to the manufacturer’s instructions. RNA samples were digested with an RNase-free DNase set (Qiagen) to remove genomic DNA and further purified using the RNeasy kit (Qiagen). Integrity of RNA samples was assessed using RNA 6000 NanoChips and an Agilent 2100 Bioanalyzer (Agilent, Santa Clara, USA). RNA samples with integrity > 7 were used further for RNA-Seq and real-time PCR (see below).

### 2.3 Transcriptomics

RNA-Seq raw reads were quality-controlled and cleaned using SOAPnuke software (http://www.seq500.com/uploadfile/SOAPnuke.zip) to remove low-quality reads, contaminated reads, and reads with adapters. The resulting clean reads were aligned to the human genome database (http://genome.ucsc.edu/) and matched with human genes using HISAT2 [31]. After counting of the reads mapped to each gene, the “fragments per kilobase per million fragments” (FPKM) method was used to normalize data in the RSEM algorithm [32] (https://deweylab.github.io/RSEM/), and low-expression genes (FPKM < 1) were filtered in each sample. NOIseq[33] was employed to calculate the log 2-fold change (log2FC) and probability value for each gene in comparison, and the following criteria were used: log2FC > 1 or < -1 and probability >0.8. Mapman software was used for PageMan analysis [34], and we considered enrichment terms to be significant if they were associated with p < 0.01.

### 2.4 Proteomics

#### 2.4.1 Label-free sample preparation

Cell pellets were suspended on ice in 200 μL lysis buffer (4% SDS, 100 mM DTT, 150 mM Tris-HCl pH 8.0). Cells were disrupted with agitation using a homogenizer (Fastprep-24®, MP Biomedical, California, USA) and boiled for 5 min. The samples were further ultra-sonicated and boiled for another 5 min. Undissolved cellular debris were removed by centrifugation at 14000 rpm for 15 min. The supernatant was collected and quantified with a BCA Protein Assay Kit (Bio-Rad, USA).

Protein (250 μg per sample) was digested according to the FASP procedure [35]. Briefly, the detergent, DTT and other low-molecular-weight components were removed using 200 μl UA buffer (8 M urea, 150 mM Tris-HCl, pH 8.0) and repeated ultrafiltration (Microcon 30 kDa, Lianke, Shanghai, China). Then 100 μL 0.05 M iodoacetamide in UA buffer was added to block reduced cysteine residues, and the samples were incubated for 20 min in the dark. The filter was washed three times with 100 μl UA buffer, then twice with 100 μl 25 mM NH_4_HCO_3_. Finally, the protein suspension was digested with 3 μg trypsin (Promega) in 40 μl 25 mM NH_4_HCO_3_ overnight at 37 °C, and the resulting peptides were collected as a filtrate. The peptide content was estimated by UV spectrometry at 280 nm assuming an extinction coefficient of 1.1 for a 0.1% (g/L) solution, which is based on the frequencies of tryptophan and tyrosine in vertebrate proteins[28].

#### 2.4.2 Liquid chromatography-tandem mass spectrometry

Each peptide sample was desalted on Empore SPE C18 cartridges (standard density, bed inner diameter 7 mm, volume 3 ml; Sigma), then concentrated by vacuum centrifugation and reconstituted in 40 µl 0.1% (v/v) trifluoroacetic acid. Peptides (5 μg) were loaded onto a reverse-phase C18 column (Thermo Scientific Easy Column, length 10 cm, inner diameter 75 μm, resin 3 μm) in buffer A (2% acetonitrile and 0.1% formic acid) and separated during 120 min with a linear gradient of buffer B (80% acetonitrile and 0.1% formic acid) at a flow rate of 250 nL/min controlled by IntelliFlow technology on an Easy nanoLC system (Proxeon Biosystems, now Thermo Fisher Scientific). The liquid chromatography system was connected to a Q Exactive electrospray ionization-tandem mass spectrometer (Waters Corporatiaon, Milford, USA). A top10 method[36, 37] was used to dynamically select the most abundant precursor ions for high energy collisional dissociation (HCD) fragmentation from the survey scan (*m/z* 300–1800). Determination of the target value was based on predictive automatic gain control, and the duration of dynamic exclusion was 25 s. Survey scans were acquired at a resolution of 70,000 at *m/z* 200, and the resolution for HCD spectra was set to 17,500 at *m/z* 200. Normalized collision energy was 30 eV, which specifies the minimum percentage of the target value likely to be reached at maximum fill time, was defined as 0.1%. The instrument was run with peptide recognition mode enabled. Mass spectrometry experiments were performed in triplicate for each sample.

#### 2.4.3 Sequence database searching and analysis

Mass spectrometry data were analyzed using freely available MaxQuant software 1.3.0.5 (www.maxquant.live) against the database uniprot_ human_152544_20160420.fasta (152,544 total entries, downloaded December 20, 2019). An initial search was set at a precursor mass window of 6 ppm. The search employed an enzymatic cleavage rule of Trypsin/P and allowed a maximum of two missed cleavage sites and a mass tolerance of 20 ppm for fragment ions. During database searching, cysteine carbamidomethylation was defined as a fixed modification, while protein N-terminal acetylation and methionine oxidation were defined as variable modifications. Label-free quantification was carried out in MaxQuant as described [38]. The peptide spectrum match was filtered by posterior error probability (PEP), which was allowed to be as high as 0.1. Peptides were included in further analysis if they showed an andromeda score > 0 and false discovery rate < 0.01. Co-fragmentation generally reduces the number of peptides identified in database searches and therefore poses problems for quantification methods based on reporter fragments, since it removes peptides that contribute to the measured ion ratios. Co-fragmentation was performed according to the default algorithm in Max-Quant software [39]. Protein abundance was calculated on the basis of the normalized spectral protein (LFQ) intensity. Our proteomics results have been deposited in the ProteomeXchange Consortium in the PRIDE repository [40] under the dataset identifier PXD006350.

#### 2.5 Docking studies of celastrol binding to predicted target proteins

Docking studies were performed with selected target proteins using YASARA [41] based on the crystal structures of the targets deposited in the RCSB Protein Data Bank (http://www.pdb.org/pdb/home/home.do). Docking simulations were performed using a CHARMM force field after adding hydrogen atoms to the proteins. The binding site was defined as residues whose side chains fell at least partly within a sphere of diameter 12 Å centered at the original ligand. Default values were used for other parameters, and genetic algorithm runs were performed for each ligand.

The technical route in this study is summarized in Figure 1.

**Figure 1.**
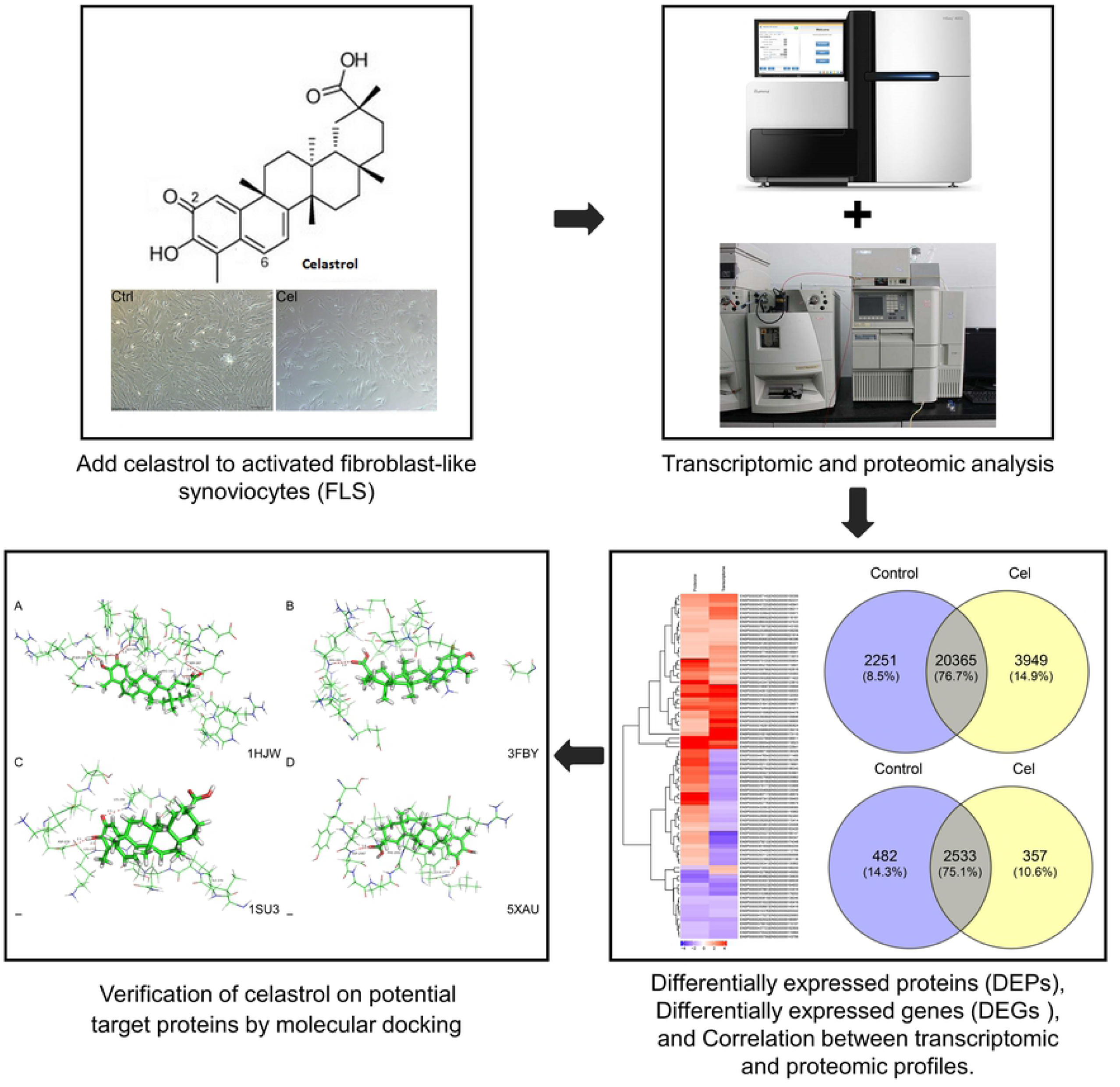
The technical route in this study.

## 3. Results

### 3.1 Identification of FLS and celastrol treatment

After confirming that the FLS expressed the marker vimentin (Figure 2A), we treated them with 5 μM celastrol for 24 h. This treatment reduced cell viability, reflected in a greater number of dead cells than in the control cultures (Figure 2B).

**Figure 2.**
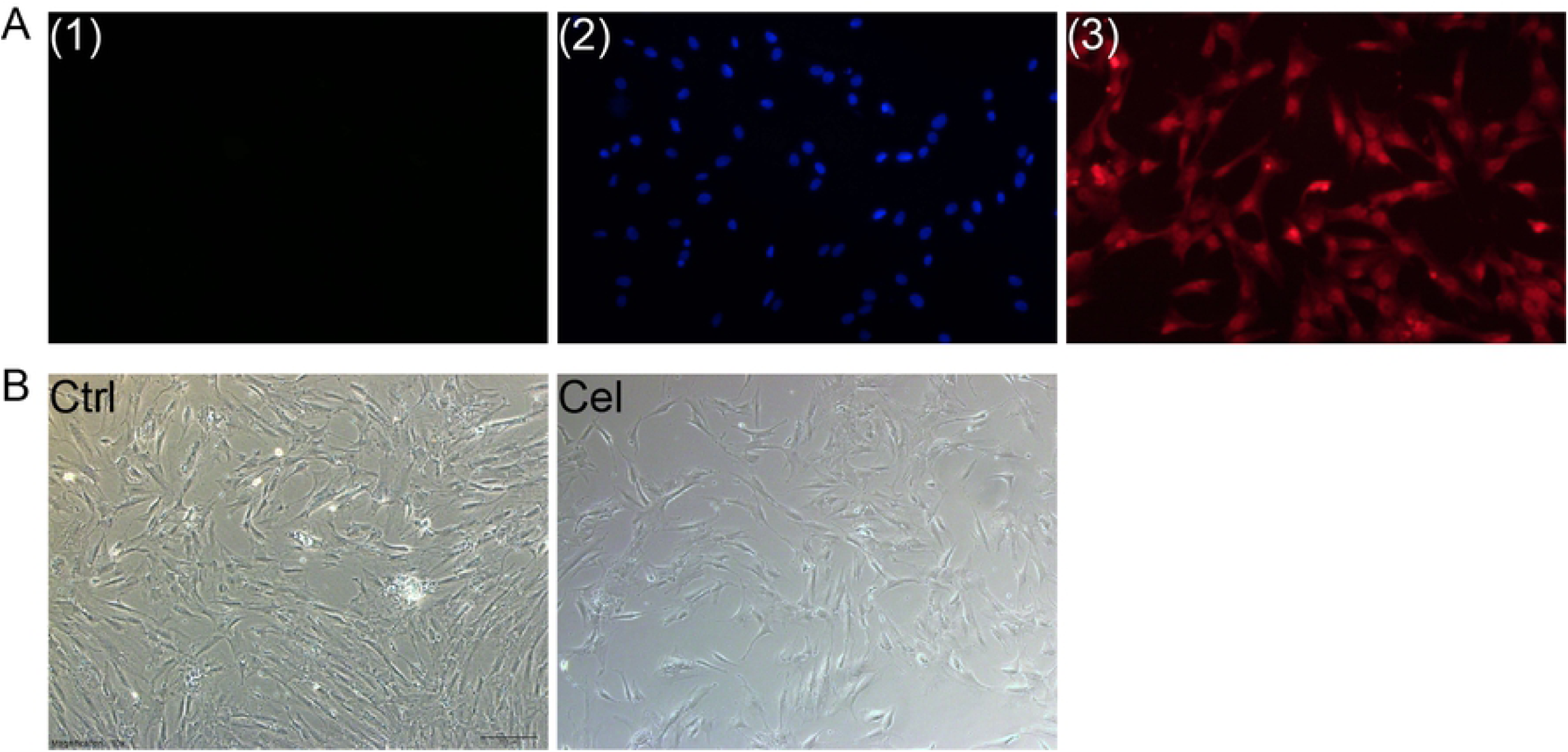
FLS culture and celastrol treatment. (A) Identification of DAPI-stained FLS cultures based on immunostaining with (1) no primary antibody, (2) anti-vimentin antibody, or (3) anti-DAPI antibody. Magnification, 200 ×. (B) Representative light micrographs of three FLS cultures treated for 24 h with DMSO vehicle (Ctrl, *upper row)* or 5 μM celastrol (Cel, *lower row*). Magnification, 10×.

### 3.2 Gene and protein expression in FLS in response to celastrol

Our analysis detected 26,565 expressed genes and 3,372 expressed proteins (in the form of 18,843 unique peptides) (Figure 3A-B). The Mascot search algorithm was used to identify proteins.

**Figure 3.**
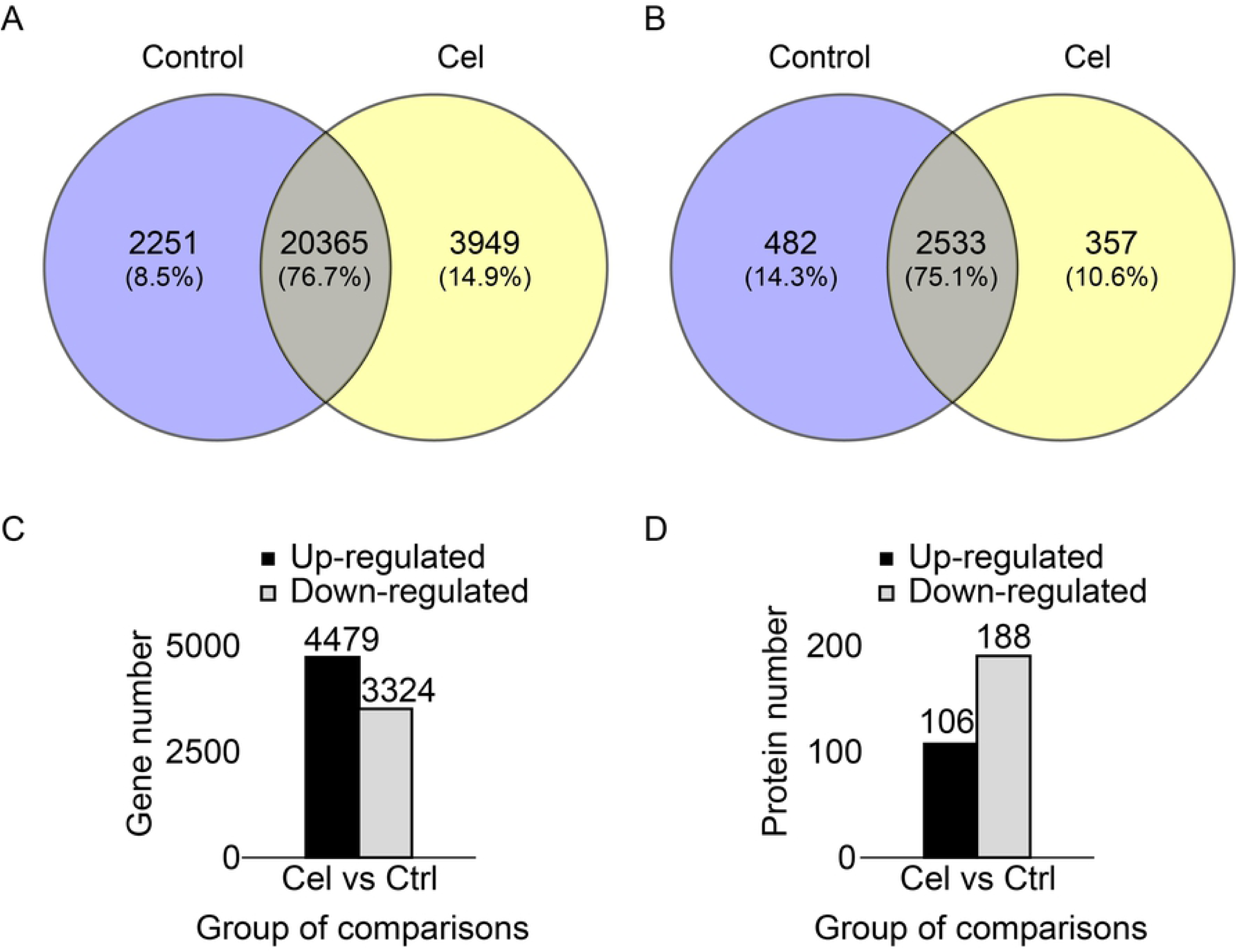
Transcriptomic and proteomic differences between FLS treated or not with celastrol. (A) Venn diagram of genes expressed in the two conditions. (B) Venn diagram of proteins expressed in the two conditions. (C) Numbers of genes differentially expressed between the two conditions. (D) Numbers of proteins differentially expressed between the two conditions.

Celastrol down-regulated 3,324 genes and up-regulated 4,479 genes, corresponding to a total of 7,803 differentially expressed genes (DEGs) (Figure 3C). The drug up-regulated 106 proteins and down-regulated 188, corresponding to a total of 294 differentially expressed proteins (DEPs) (Figure 3D).

### 3.3 Gene ontology of DEGs and DEPs

The distribution of DEPs and DEGs was analyzed across the three gene ontology categories of biological processes, cellular components, and molecular functions (http://geneontology.org). Among biological processes, DEGs and DEPs were enriched in processes related to response to oxidative stress, response to oxygen levels, extracellular structure and regulation of small-molecule metabolic processes. Among cellular components, DEGs and DEPs were enriched in collagen-containing extracellular matrix, focal adhesion, cell-substrate adherens junction and cell-substrate junction (Figure 4A). Among molecular functions, DEGs and DEPs were enriched in binding of cell adhesion molecules, ubiquitin-like protein transferase activity and ubiquitin-like protein transferase activity followed by structural components of extracellular matrix.

**Figure 4.**
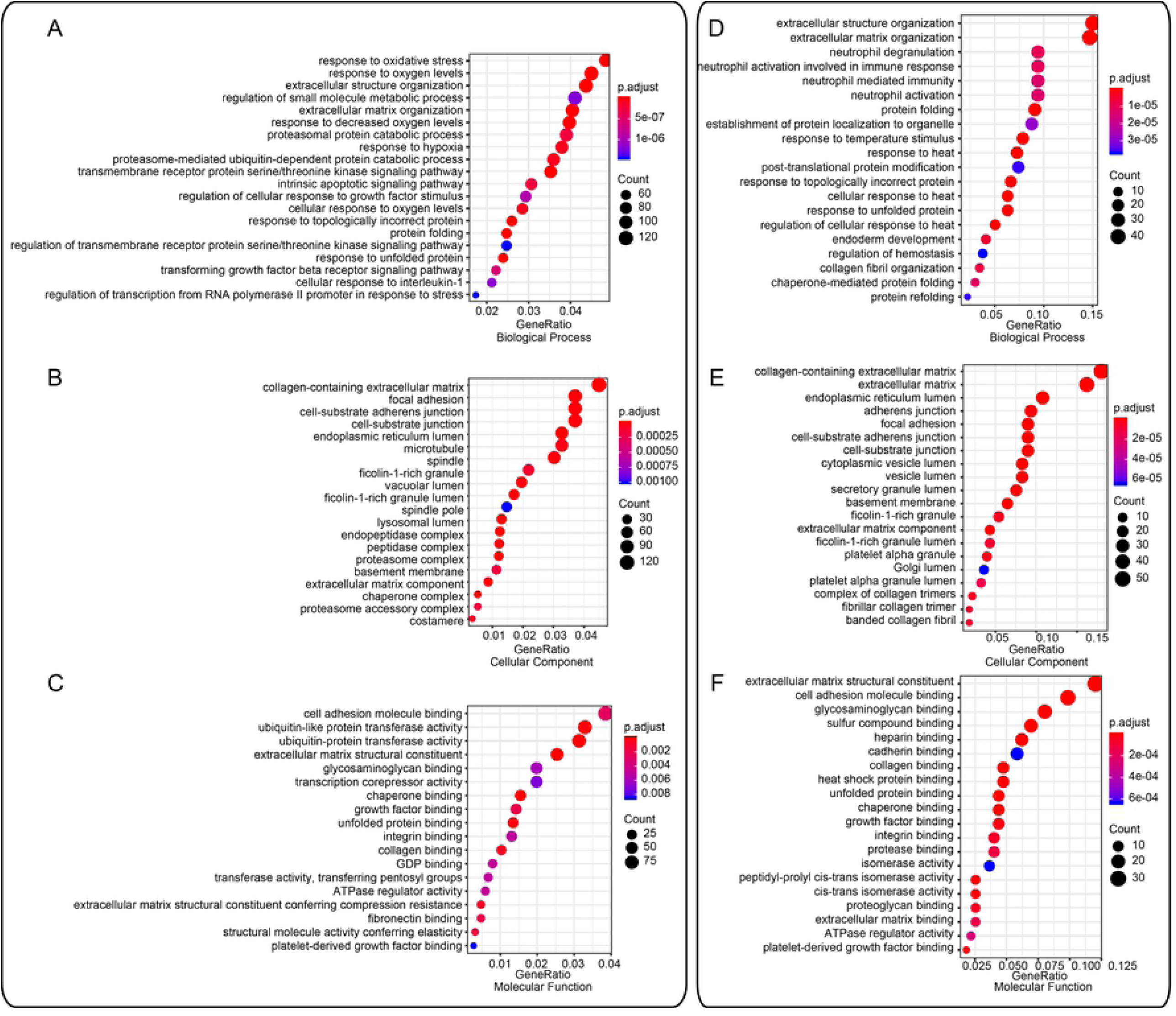
Distribution of differentially expressed genes and proteins across the three gene ontology categories of biological processes, cellular components, and molecular functions. (A-C) ClusterProfiler analysis of the transcriptome. (D-F) ClusterProfiler analysis of the proteome. The color scales indicate the different thresholds of adjusted p-values, and the sizes of the dots represent the numbers of genes assigned to the given term. The panels in this figure were generated on the website http://metascape.org/gp/index.html.

### 3.4 Correlation between proteomic and transcriptomic analyses

We observed a weak correlation between levels of transcripts and levels of their cognate proteins when we examined the entire set of genes and proteins (r = 0.2067), as well as the subset of DEGs and DEPs (r =0.2507). Altogether, we identified 36 genes up-regulated and 15 genes down-regulated at both the RNA and protein levels in response to celastrol. We identified another 27 genes that were transcriptionally down-regulated but translationally up-regulated, and two genes that were transcriptionally up-regulated but translationally down-regulated (Figure 5).

**Figure 5.**
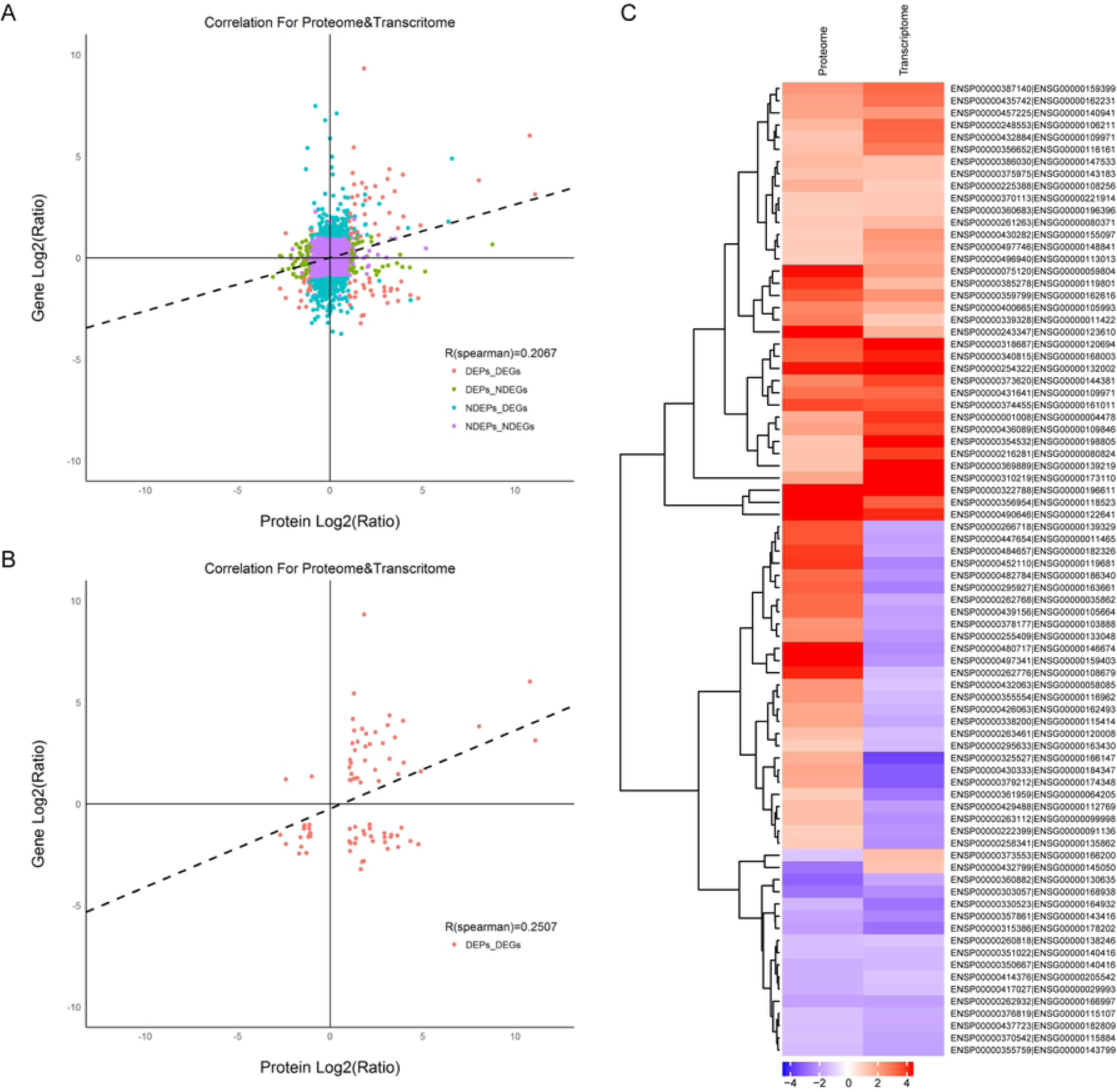
Correlation between transcriptomic and proteomic profiles in the presence and absence of celastrol. (A) Correlation analysis across all genes and proteins in the two conditions. (B) Correlation analysis of only differentially expressed genes (DEGs) and differentially expressed proteins (DEPs). (C) Hierarchical clustering of DEGs and DEPs. Red color denotes up-regulation; blue, down-regulation.

### 3.5 Docking of celastrol in target proteins

Our transcriptomic and proteomic analyses suggested that celastrol acts against several proteins involved in the extracellular matrix of FLS. Therefore, we explored the potential binding of celastrol to several such proteins whose crystal structures are known (Figure 6). In all cases, we managed to obtain reasonable complexes in which the primary forms of interaction are hydrogen bonding and hydrophobic interactions. For instance, celastrol is predicted to interact with CHI3L1 via hydrogen-bonds with Ser103, Gly143, Ser187, and Arg144 (Figure 6A). The drug is predicted to interact with ANXA6 via hydrogen-bonds with Lys224 and hydrophobic interaction with Glu643 (Figure 6B).

Fig 6

**Figure 6.**
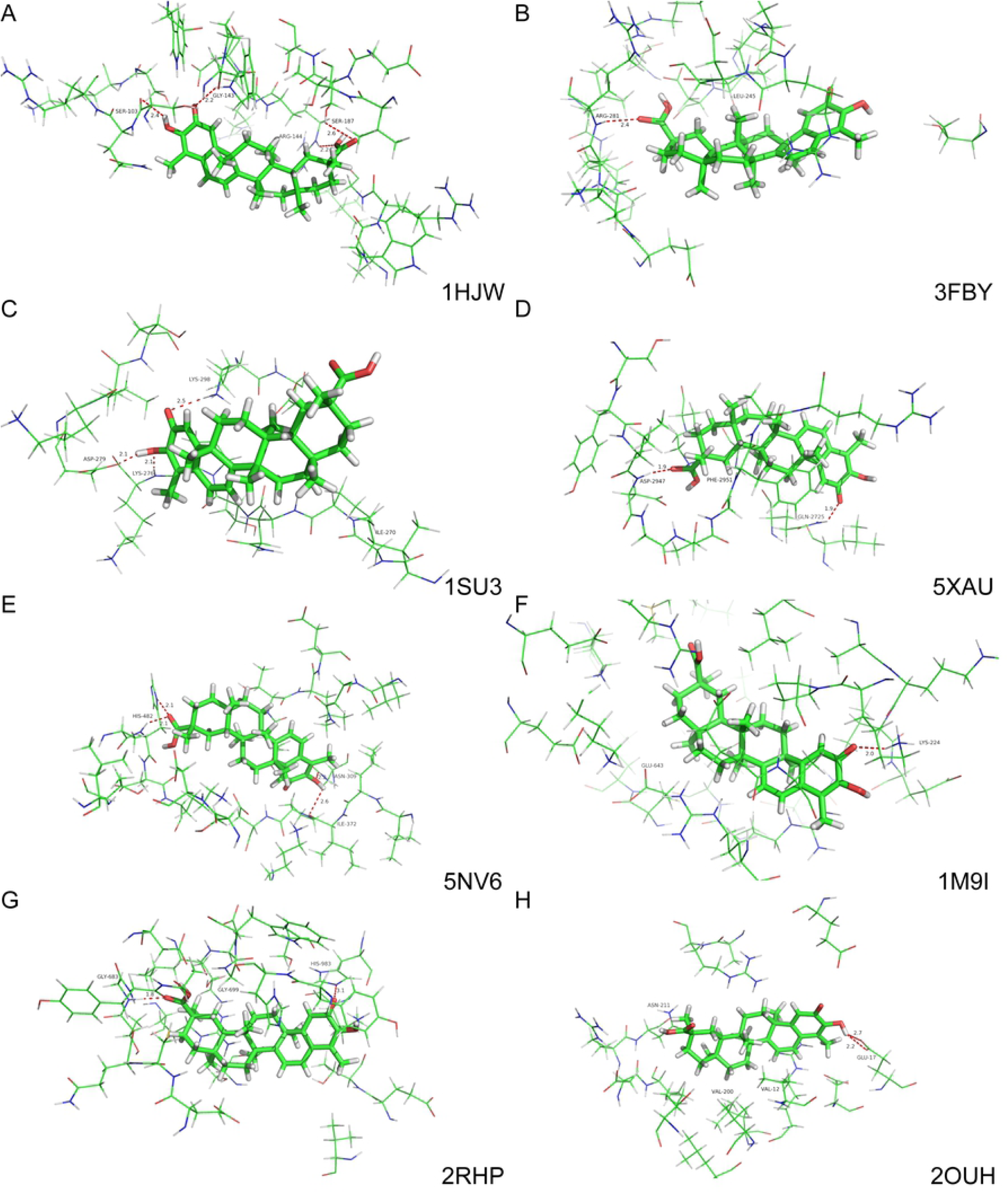
Docking studies of celastrol with potential target proteins involved in the extracellular matrix of FLS. Ball-and-stick diagrams are shown for the region within 12 Å of 8 proteins. Key protein residues are labeled. Hydrogen bonds are shown as dashed lines, together with interaction distances in Å.

(A) Chitinase 3 like 1 (PDB ID: 1HJW)[42].

(B) Cartilage oligomeric matrix protein (3FBY)[43].

(C) Matrix metalloprotease 1 (1SU3)[44]

(D) Laminin subunit gamma 1 (5XAU)[45]

(E) Transforming growth factor beta-induced (5NV6)[46]

(F) Annexin A6 (1M9I)[47]

(G) Thrombospondin 2 (2RHP)[48]

(H) Thrombospondin 1 (2OUH)[49].

## 4. Discussion

Reflecting the fact that RA is an inflammatory disease, markers of inflammation such as C-reactive protein (CRP), IL-6, and TNF-α are highly expressed in synovial fluid and serum of arthritic patients, and the levels of these markers in blood correlate with disease severity. In RA, the primary site of inflammation is the synovial tissue, from which cytokines may be released into the systemic circulation. These cytokines are involved in almost all aspects of articular inflammation and destruction in RA[50, 51].

Our transcriptomic and proteomic study of the effects of celastrol on FLS suggests that the drug exerts anti-inflammatory effects by affecting response to oxidative stress, response to oxygen levels, extracellular organization and regulation of small-molecule metabolism. Celastrol may interfere directly with the extracellular matrix and inhibit FLS proliferation and survival.

We found only modest concordance between the set of transcripts and the set of proteins expressed in FLS. This is consistent with transcriptomic and proteomic studies in lung adenocarcinomas and embryonic stem cells [52, 53]. This likely reflects, at least in part, the translational regulation and post-translational processes that affect the proteome [54].

In RA, hypoxia is a prominent feature of the micro-environment, which arises through increased oxygen consumption by inflamed resident cells and infiltrating immune cells, and through disruption of the blood supply due to vascular dysfunction. Hypoxia triggers several signaling pathways that drive RA pathology. The primary signaling pathway activated by hypoxia involves hypoxia-inducible factors (HIFs), which are highly expressed in the synovium of RA. HIFs induce inflammation, angiogenesis, cell migration, and cartilage destruction, and they inhibit the apoptosis of synovial cells and inflammatory cells. HIF expression, like that of inflammatory cytokines, can be triggered through pathways dependent and independent of oxygen, aggravating the disease[55-58]. Indeed, the number of HIF-1α-positive cells is directly related to the extent of infiltration by inflammatory endothelial cells and to the severity of synovitis [59, 60]. Excessive production of reactive oxygen species (ROS) causes the breakdown of proteoglycans, hyaluronic acid, chondroitin sulfate, and collagen [61], which destroys the joints. Our results showed that celastrol may reduce HIF expression, kill FLS and alleviate inflammation in RA.

Our results further suggest that celastrol may target several proteins in the extracellular matrix. This suggestion is based on transcriptomic and proteomic profiling, as well as docking simulations. Drugs that target the extracellular matrix may have advantages over drugs that target proteins in the synovium because most drugs are rapidly removed from the synovium[62, 63], whereas matrix proteins experience slow turnover, prolonging the therapeutic window.

## 5. Conclusions

Our combination approach has generated several testable hypotheses about proteins and cellular pathways through which celastrol may reduce inflammation in RA. This work provides a wide-reaching foundation for further genetic and biochemical studies to improve our understanding of RA and its treatment.

## Abbreviations

RA: rheumatoid arthritis
FLS: fibroblast-like synoviocytes
MMP: matrix metalloprotease
HCD: high energy collisional dissociation
DEPs: expressed proteins
DEGs: differentially expressed genes
CRP: C-reactive protein
HIFs: hypoxia-inducible factors
ROS: reactive oxygen species

## Conflicts of interest

The authors have no conflicts of interest to declare.

### Acknowledgments

This study was supported by the National Natural Science Foundation of China (grant no. U1804179), the Henan Science and Technology Innovation Team, the Investigation on Plant Resources in Dabie Mountains and the Study and Utilization of Active Components of Special plants (grant no. 2017083), Henan key scientific and technological projects (202102310190) and the Nanhu Scholars Program for Young Scholars of Xinyang Normal University (grant no. 2018001).

## Declarations

1 The manuscript doesn’t contain human participants, human data or human tissue;

2 All the authors claim that the article has not been published and is not under consideration for publication elsewhere and all the authors have approved and agreed to bear the applicable publication charges if their manuscript is accepted for publication.

3 Availability of data can be downloaded from attachments

4 Competing interests

The authors have no conflicts of interest to declare.

## 6 Authors’ contributions

Conceived and designed the experiments: Xinqiang Song.

Performed the experiments: Erqin Dai.

Analyzed the data: Yu Zhang, Lei Wang, Hongtao Du

Wrote the paper: Xinqiang Song.

